# From fruit growth to ripening in plantain: a careful balance between carbohydrate synthesis and breakdown

**DOI:** 10.1101/2021.12.03.471126

**Authors:** N.A. Campos, S. Colombie, Annick Moing, C. Cassan, D. Amah, R. Swennen, Y. Gibon, S.C. Carpentier

**Author notes:** Correspondence author: Dr. Sebastien Carpentier, Department of Biosystems, KULeuven, Willem Decroylaan 42 bus 2455- 3001 Leuven, Belgium. S.C.C. and R.S. conceived the original screening and research plans; S.C. and Y.G. supervised the experiments; N.C. performed the experiments using the proteomics methods and S.C and A.M. performed the experiments based on the metabolome methods; D.A. performed the field experiment; C.C. provided technical assistance; S.C.C, N.C, S.C, A.M analyzed the data; N.C, and S.C.C. wrote the article with contributions of all the authors; S.C.C. supervised and completed the writing. S.C.C. agrees to serve as the author responsible for contact and ensures communication. We acknowledge USAID for the project AID-BFS-G-II-00002-11 Reviving the plantain breeding program at IITA – International Institute for Tropical Agriculture. The authors would furthermore like to thank all donors who supported this work through their contributions to the CGIAR Fund (http://www.cgiar.org/who-we-are/cgiar-fund/fund-donors-2/), and in particular to the CGIAR Research Program on Roots, Tubers and Bananas, and the PHENOME (French ANR-11-INBS-0012) project for funding. The metabolite analyses were performed on Bordeaux Metabolome facility.

## Abstract

We investigated the fruit development in two plantain banana cultivars from two weeks after bunch emergence till twelve weeks through high-throughput proteomics, major metabolite quantification and metabolic flux analyses. We give for the first time an insight at early stages of starch synthesis and breakdown. Starch and sugar synthesis and breakdown are processes that take place simultaneously. During the first eight to ten weeks the balance between synthesis and breakdown is clearly in favour of sugar breakdown and a net starch synthesis occurs. During this period, plantain fruit accumulates up to 48% of starch. The initiation of the ripening process is accompanied with a shift in balance towards net starch breakdown. The key enzymes related to this are phosphoglucan water dikinase (PWD), phosphoglucan phosphatase, α-1,6-glucosidase starch debranching enzyme (DBE), alpha glucan phosphorylase (PHS) and 4-alpha glucanotransferase disproportioning enzyme (DPE). The highest correlations with sucrose have been observed for PHS and DPE. There is also a significant correlation between the enzymes involved in ethylene biosynthesis, starch breakdown, pulp softening and ascorbate biosynthesis. The faster ending of maturation and starting of ripening in the Agbagba cultivar are linked to the key enzymes 1-aminocyclopropane-1-carboxylate oxidase and DPE. This knowledge of the mechanisms that regulate starch and sugar metabolisms during maturation and ripening is fundamental to determine the harvest moment, reduce postharvest losses and improve final product quality of breeding programs.

## 1 Introduction

Fruit development is a complex phenomenon that encompasses several overlapping stages: cell division, cell enlargement, maturation, ripening and senescence (Paul et al., 2012). At the initiation of fruit development, fruits enlarge mainly through cell division and they reach their final size by increasing cell volume. The active cell division and cell expansion are accompanied by a net accumulation of storage products until full maturation. Ripening induces changes in flavour, texture, colour, and aroma. Fruits can be divided into two groups with contrasting ripening mechanisms. Climacteric fruit (such as tomato, avocado, apple, and banana) are linked to ethylene biosynthesis and an increase in respiration which induces ripening (White, 2002). During maturation, two systems of ethylene are operational: system 1 and system 2. During early maturation, system 1 is active and the rate of ethylene production is basal and there is an auto-inhibition of ethylene production. But as maturation progresses, this inhibition process is stopped and there is an auto-induction of ethylene production leading to the onset of ripening (Paul et al., 2012).

Edible bananas are parthenocarpic and so the ovaries develop into seedless fruits without pollination stimulus. The pulp-initiating cells are situated within the inner epidermis of the fruit pericarp and septa. In parthenocarpic bananas, those cells start to proliferate very fast after flowering (bunch emergence) (Ram et al., 1962). The increase in cell number in the initiating region of the pulp continues up to about 4 Weeks After bunch Emergence (WAE). Then it subsides and growth is largely realized by cell enlargement (Ram et al., 1962). Sugar deposition and starch synthesis in the pulp cells commence very early and they become well established by 8 WAE. The first signs of starch disappearance have been reported to be around 12 WAE (Ram et al., 1962). However, this is dependent on the environment and on the genotype. Depending on the genotype, banana fruit has been reported to accumulate between 12 and 35% of starch during 4-8 WAE and from 8 WAE starch content drops to between 15 and 0% in late stages of maturation (Soares et al., 2011; Cordenunsi-Lysenko et al., 2019). Plantains are part of the group of bananas that accumulate a large amount of starch. At ripe stage, plantains still have a high starch content, which affects their taste (Soares et al., 2011). Therefore, plantains are not suitable as sweet dessert bananas and are consumed as starch source. The current practice is to harvest when fruits of the first hand show signs of ripening (Dadzie and Orchard, 1997). Plantains are an important staple food in tropical and subtropical countries, being of special importance in West-Africa (Vuylsteke et al., 1993). Genetically, they are triploids and belong to the AAB genotype group. They are a product of a natural cross between *Musa acuminata* (A genotype) and *Musa balbisiana* (B genotype) (Simmonds, 1962). And although morphologically they are quite diverse, genetically they are extremely uniform (Crouch et al., 2000). The recent release of the B genome suggested a dominance of genes related to starch metabolism, leading to a higher starch accumulation during fruit development (Wang et al., 2019). A better understanding of the mechanisms that regulate sugar primary metabolism during fruit development will be important to select hybrids with the best post-harvest traits.

Most papers are focused on Cavendish sweet banana that is the most exported banana cultivar in the world (Agopian et al., 2008; Toledo et al., 2012; Asif et al., 2014; Du et al., 2016). Recently, we published the first proteome of plantain fruit and a comparison of the proteomes of Cavendish and plantain during the final ripening process (Campos et al., 2018; Bhuiyan et al., 2020). Together with the recent update from the B genome sequence (Wang et al., 2019), these works are contributing to elucidate the fruit development in plantain as well to determine the role of the B genome in fruit quality. In complement to proteome studies, metabolic flux can be predicted using constraint-based models based on metabolic network description through stoichiometric equations of reactions, and on the assumption of pseudo-steady state and the choice of an objective function (Orth et al., 2010). Such knowledge-based stoichiometric models describing central metabolism have already proved usefull in tomato fruit to estimate fluxes throughout the development and to show that carbon degraded from starch and cell wall generated an excess of energy dissipated just before the onset of ripening coinciding with the respiration climacteric (Colombié et al., 2015; Colombié et al., 2017). By combining proteomics and flux studies, we gain here unique insights into the order of appearance and dominance of specific enzymes/fluxes involved in starch synthesis and breakdown and sugar synthesis in plaintain fruit.

## 2 Material and Methods

### 2.1 Biological Material

The biological samples were collected from the IITA Experimental Field in Ibadan, Nigeria, during the period from October 2016 to February 2017.

Five banana plants from Agbagba and Obino L’Ewai cultivars were selected and the same plants were followed during all the experiment. One fruit per plant was collected from 2 WAE until the fruits reached full maturity. The collected fruits were cleaned and measurements of fruit length (L) and circumference (C) were taken. For the fruit volume calculation our formula was based on (Simmonds, 1953). To know the correlation between fruit weight, calculated fruit volume and real fruit volume, the real volume of representative fruits was measured by submerging the them in water in a measuring cylinder. For the remainder of the fruits, the volume was calculated with the formula: Volume (cm^3^) = ((Fruit length * (Fruit circumference)^2^ *0.0616)+0.3537). After determination of fruit length and circumference, the fruit was separated from the peels, cut in smaller pieces and stored at – 80 °C until lyophilization. Samples were lyophilized to ensure a safe transportation from Nigeria to Belgium and to facilitate the protein and metabolite extraction process (Carpentier et al., 2007). The lyophilized samples were then, sent to Belgium where the protein extraction, quantification and identification were performed and to France for metabolite analysis.

### 2.2 Protein extraction, quantification, identification and annotation

Extractions were performed following the phenol-extraction/ammonium-acetate precipitation protocol described previously (Carpentier et al., 2005; Buts et al., 2014). Samples of 2 and 4 WAE could not be analyzed through proteomics due to the presence of many interfering compounds disturbing the correct application of the protocol.

After extraction, 20 µg of proteins were digested with trypsin (Trypsin Protease, MS Grade ThermoScientific, Merelbeke, Belgium) and purified by Pierce C18 Spin Columns (ThermoScientific, Merelbeke, Belgium). The digested samples (0.5µg/5µL) were separated in an Ultimate 3000 (ThermoScientific) UPLC system and then in a Q Exactive Orbitrap mass spectrometer (ThermoScientific) as described (van Wesemael et al., 2018). For protein quantification, we used the software Progenesis® (Nonlinear Dynamics). In this software we used MASCOT version 2.2.06 (Matrix Science) against the Musa V2 database of *M. acuminata* and *M. balbisiana* (Martin et al., 2016; Wang et al., 2019) (157832 proteins). Tandem mass spectra were extracted by Progensis. All MS/MS spectra were searched with a fragment ion mass tolerance of 0,02 Da and a parent ion tolerance of 10 PPM. Carbamidomethyl of cysteine was specified in Mascot as a fixed modification. Deamidation of asparagine and glutamine and oxidation of methionine were specified in Mascot as variable modifications and the results were reintroduced in Progenesis. Scaffold (version Scaffold_4.11.0, Proteome Software) was used to validate MS/MS based peptide and protein identifications. Peptide identifications were accepted if they could be established at greater than 95,0% probability by the Peptide Prophet algorithm with Scaffold delta-mass correction (Keller et al., 2002; Searle, 2010). Protein identifications were accepted if they contained at least 1 identified peptide. Protein probabilities were assigned by the Protein Prophet algorithm (Nesvizhskii et al., 2003). Proteins that contained similar peptides and could not be differentiated based on MS/MS analysis alone were grouped to satisfy the principles of parsimony. Proteins sharing significant peptide evidence were grouped into clusters. A protein false discovery rate of 0.8% and a spectral false discovery rate of 0.04% was observed by searching the reverse concatenated decoy database (157832 proteins). All data have been made available in the public repository PRIDE under the project name: Carbohydrate metabolism during plantain development, Project accession: PXD029901 and Project DOI: 10.6019/PXD029901.

Gene annotations were taken from the banana Hub (Droc et al., 2013) and verified in Plant Metabolic Network (Hawkins et al., 2021) and Prosite (Expasy SIB Bioinformatics Resource Portal). Subcellular prediction were analyzed via the software DeepLoc 1.0 (Almagro Armenteros et al., 2017).

### 2.3 Metabolic Analysis

To complement our proteomics data and improve our insights about plantain fruit development, we analyzed major metabolic traits in pulp. Metabolites were extracted from 10 mg aliquots of lyophilized ground samples via three successive extractions with ethanol-buffer mixtures successively composed of 80, 80 and 50% ethanol and 10 mM Hepes/KOH buffer (pH 6). The supernatants were collected and pooled in order to measure soluble metabolites. Glucose, fructose and sucrose were measured enzymatically (Stitt et al., 1989). Glucose-6-phosphate, fructose-6-phosphate and glucose-1-phosphate were measured using an enzyme cycling assay (Gibon et al., 2002). Malate was measured enzymatically as in (Mollering, 1985). Total free amino acids were measured using fluorescamine (Bantan-Polak et al., 2001). Polyphenols were measured using Folin-Ciocalteu’s reagent (Blainski et al., 2013). In order to quantify the total protein content, the pellets were resuspended in 100 mM NaOH and then heated for 20 min. After centrifugation (5,000 *g*, 5 min), the total protein content was measured with Coomassie Blue (Bradford, 1976). After neutralization with HCl, starch was quantified in the pellets as described previously (Hendriks et al., 2003). Finally, the pellet was washed twice with water and twice with ethanol 96% v/v, dried and weighed to estimate the cell wall content.

### 2.4 Flux calculation by constrain-based modelling

A flux-balance model was constructed by integrating biochemical and physiological knowledge about central metabolism previously described (Colombié et al., 2015; Soubeyrand et al., 2018) dedicated to breakdown and transformation of extracellular nutriments to produce energy and metabolites and a specialized metabolic pathway producing the main polyphenol compounds. Energy intermediates, both ATP and NAD(P)H, were explicitly considered and all the cofactors were defined as internal metabolites, which means that they were balanced, thus constraining the metabolic network not only through the carbon and nitrogen balance but also through the redox and energy status.

To solve the flux balance model, constraints were applied for (1) flux reversibility or irreversibility and for (2) outfluxes boundaries. Therefore, concentrations of accumulated metabolites and biomass components, expressed on a mole per fruit basis, were fitted to calculate the corresponding fluxes. Stoichiometric network reconstruction encompassing central and polyphenol metabolism and mathematical problems were implemented using MATLAB (Mathworks R2012b, Natick, MA, USA) and the optimization toolbox, solver quadprod with interior-point-convex algorithm for the minimization. Flux maps were drawn with the flux visualization tool of VANTED 2.1.0.

### 2.5 Statistical analyses

For proteins, statistical analyses were made using the software Statistica 8 (TIBCO) based on the exported protein quantifications of Progenesis. We performed a principal component analysis (PCA, with NIPALS algorithm) to get an overview of the proteome data. We performed a partial least squares analysis (PLS) (NIPALS algorithm) to differentiate proteins with a significant correlation to the time points, the genotype, metabolite using all protein quantifications as continuous predictors (x matrix) and the time points, genotype and quantified metabolite as dependent variables (y matrix). We applied a two-way ANOVA (*P*<0.05) to the selected proteins to verify their significance affected by the time point, genotype or the interaction between both.

For metabolites, a principal component analysis was performed on the averages per cultivar and time point.

All displayed regressions were made in Microsoft excel and based on the best fit R^2^. Pearson correlations between proteins or between proteins and selected metabolites or other variables were calculated with Statistica 8 (TIBCO).

To integrate the different omics data, the protein inference and isoform redundancy issue was tackled by quantifying the proteins at protein cluster level and EMPAI quantification (Scaffold_4.11.0, Proteome Software). To find the protein clusters that correlated to the modelled fluxes, we performed a two-block sparse partial-least-squares discriminant analysis (sPLS-DA) with mixOmics package of R (Rohart et al., 2017) using DIABLO application (Singh et al., 2019) with default parameters. To the relationships between the proteins and fluxes were calculated with *P*<0.001 (after false discovery rate [FDR] correction) threshold for Pearson correlations.

## 3 Results

### 3.1 The different growth stages are characterized by a particular proteome and metabolic profile

Based on an unsupervised principal component analysis, the proteome differed at each time point (Figure 1). The first component explained 23% of total variability and clearly separated the different time points. The second component explained 17% of total variability, and was correlated to the cultivar. Both cultivars had a similar proteome except for the last time point at 12 Weeks After bunch Emergence (WAE). The same is true for the metabolite and the flux analysis data, expect that the largest difference between the two cultivars is observed at 6 WAE for the metabolites and both at 6 and 12 WAE for the fluxes (Figure S1).

**FIGURE 1:**
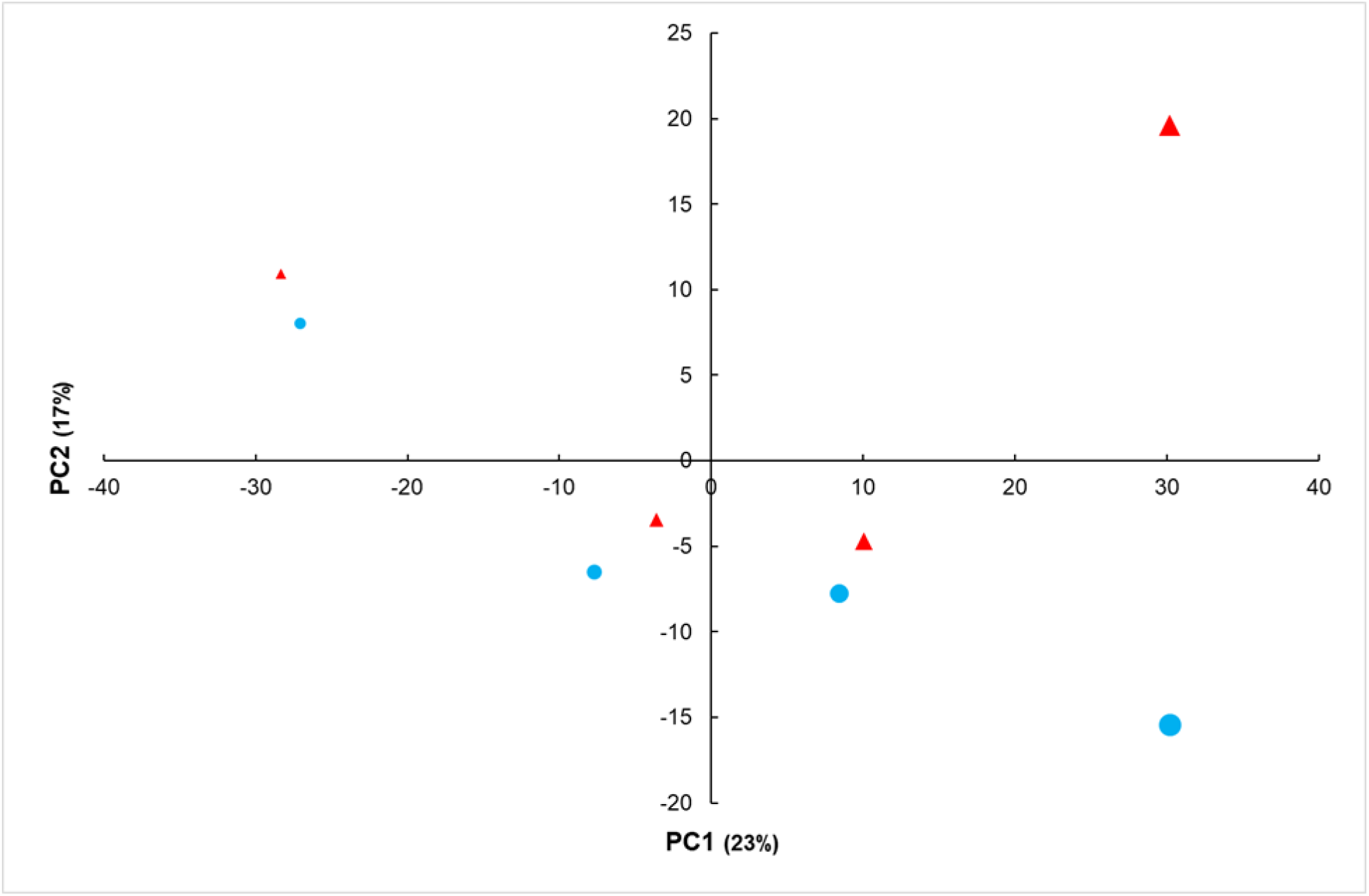
Principal Component Analysis of the proteomics data (2183 proteins) of the two varieties of plantain banana during fruit developement. Displayed are the average scores per cultivar and time point. Agbagba (blue) and Obino l’Ewai (red). The size of the data points is proportional to the time of sampling. Pulp samples were analyzed at 6, 8, 10 and 12 WAE, n =3-5.

Concerning fruit growth, in both cultivars we observed a sigmoid curve with three growth phases: a fast growth phase (0-6 WAE), a phase of slow growth (6-8 WAE) and a second phase of fast growth (8-12 WAE) (Figure 2, S2). Based on the observed abundance pattern of Sucrose Synthase (SuSy) and invertase, we hypothesize that the first fast growth phase is completely dominated by Sucrose Synthase (SuSy), while the third growth phase is dominated by invertase. The abundance of invertase showed an excellent correlation with the growth rate (Figure 3).

**FIGURE 2:**
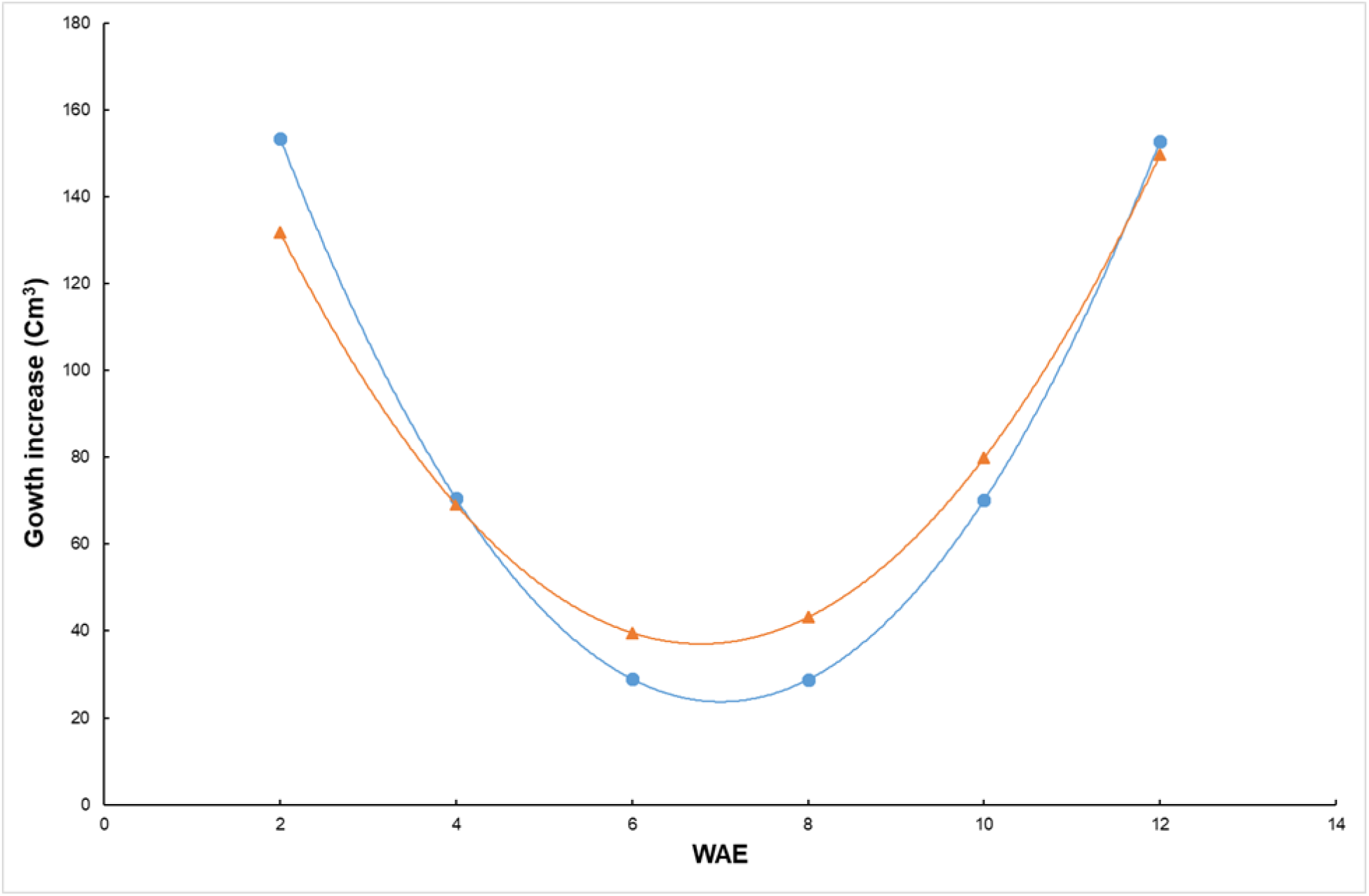
Changes in growth of fruit of two plantain varieties from 2 to 12 WAE (derivative of cubic regression model). AG: Agbaba (Blue); OB: Obino l’ewai (Red).

**FIGURE 3:**
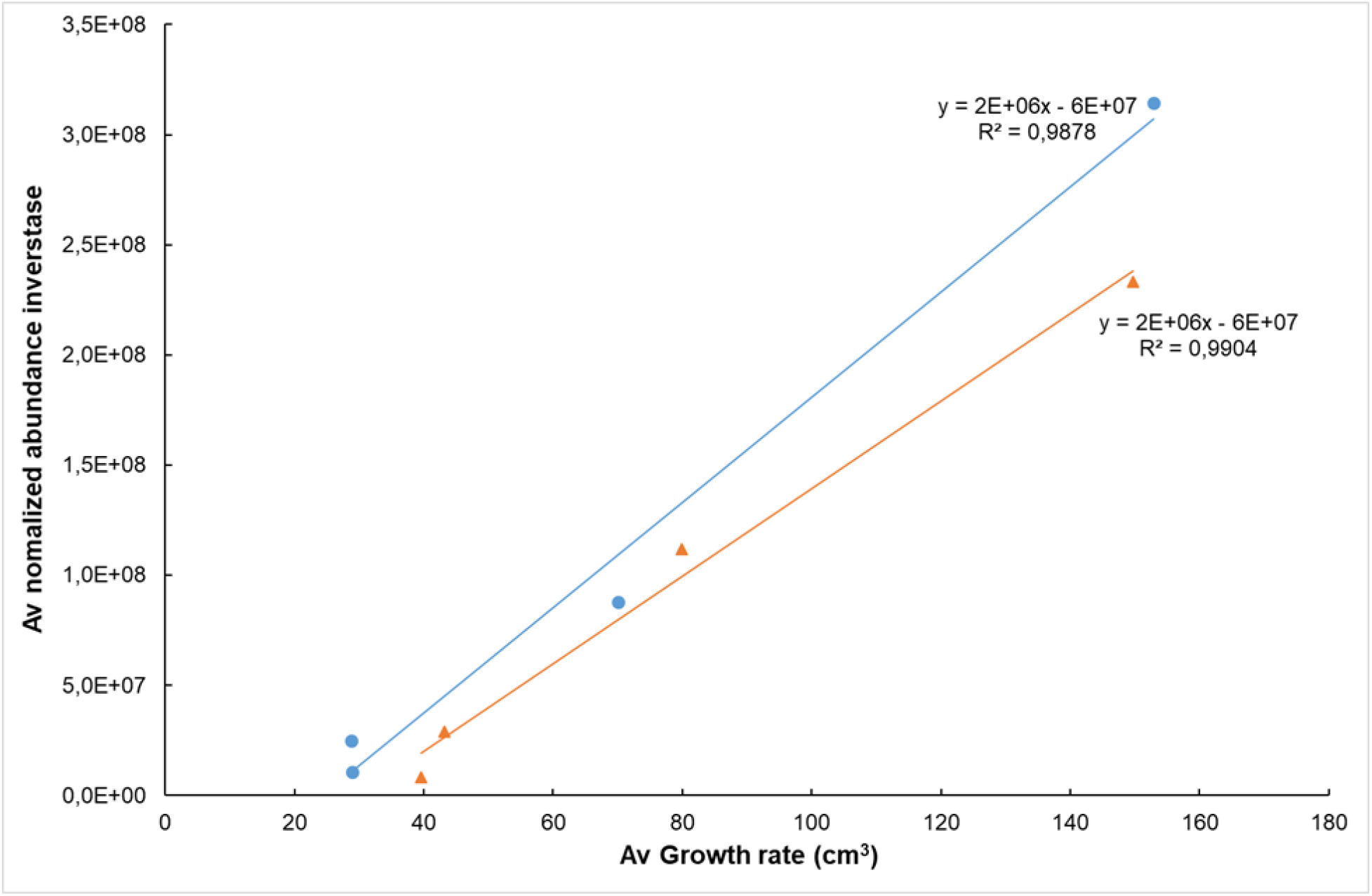
Correlation between the average growth rate (6-12 WAE) and the average normalized abundance of cytoplasmic invertase Mba10_g13890.1 for two plantain varieties. Agbaba (blue); Obino l’ewai (red).

We did find evidence to confirm the involvement of cell division in growth in our proteomics data. Based on the identified histone proteins we deduce that cell division takes place up till 8 WAE (Table 1). A fast cell division is also accompanied with a high activity of cell wall building and modifying enzymes (UDP-glucose 6-dehydrogenase, UDP-glucuronic acid decarboxylase, Beta-glucosidase), mRNA translation (eukaryotic initiation factors, ribosomal proteins), protein folding (T-complex proteins) and turnover (proteasome complex) (Table 1). The identified proteins involved in the cell division processes significantly decrease in abundance from 6 WAE.*3.2 Starch and sugar metabolism: synthesis and breakdown are processes that take place simultaneously*

**Table 1:**
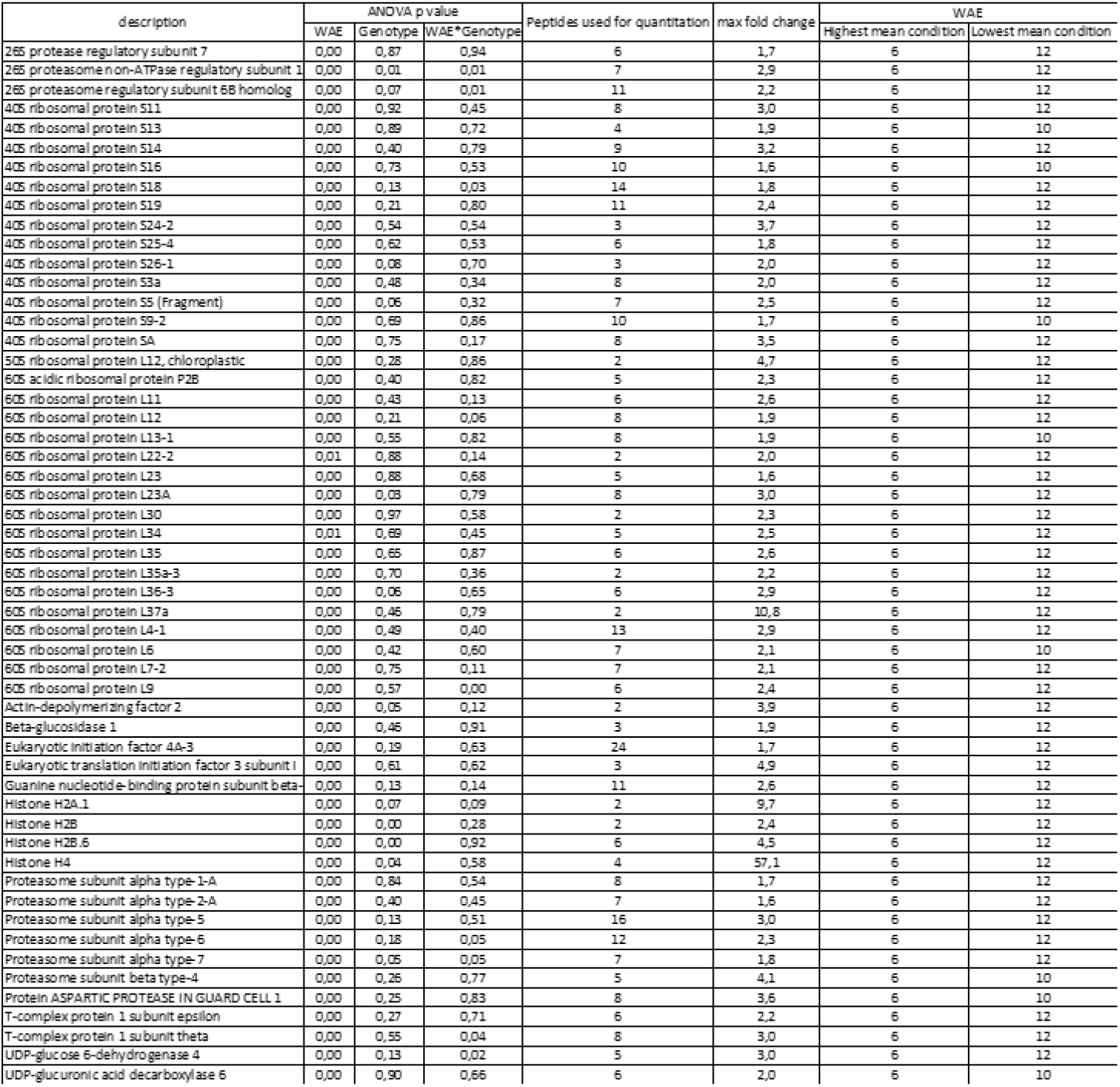
Proteins linked to growth in plantain banana pulp WAE: Weeks After Emergence This table only displays the ANOVA p-values of the protein paralog with the lowest value. The full list of significant protein paralogs can be seen in Table S1.

The pulp at 6 WAE contained three times more fructose than glucose, but the concentration of fructose represented <1 % of that of starch and less than 15% of that of sucrose. Among the hexose phosphates, the amount of Glc-6-P was 20-fold higher than Glc-l-P (Table 2). The pulp at 12 WAE contained twice more fructose than glucose, but the concentration of fructose represented <0.5 % of that of starch and less than 5% of that of sucrose. Among the hexose phosphates, the amount of Glc-6-P was 20-fold higher than Glc-l-P (Table 2).

**Table 2:** Metabolite data of plantain banana pulp. Homogeneous groups over time within the same metabolite are indicated by a letter. A<B<C<D<E, Groups sharing a letter are not significantly different (Fisher test). WAE: weeks after emergence OB: Obino ‘l Ewai AG: Agbagba

|  |  |  |  |  |  |  |  |  |  |  |  |  |
| --- | --- | --- | --- | --- | --- | --- | --- | --- | --- | --- | --- | --- |
| time point<br>WAE | 2 | 4 | 6 | 8 | 10 | 12 | 2 | 4 | 6 | 8 | 10 | 12 |
| Genotype | AG | AG | AG | AG | AG | AG | OL | OL | OL | OL | OL | OL |
| Biological<br>replicates | 5 | 5 | 5 | 5 | 5 | 4 | 5 | 5 | 5 | 5 | 5 | 4 |
| | Mean $\mu\text{mol/gDW}$ | | | | | | | | | | | |
| F6P | 0,5<sup>D</sup> | 0,4<sup>ABCD</sup> | 0,3<sup>AB</sup> | 0,3<sup>AB</sup> | 0,3<sup>ABC</sup> | 0,5<sup>CD</sup> | 0,5<sup>CD</sup> | 0,4<sup>ABCD</sup> | 0,3<sup>A</sup> | 0,3<sup>AB</sup> | 0,4<sup>BCD</sup> | 0,4<sup>ABCD</sup> |
| Fructose | 82,7<sup>C</sup> | 51,5<sup>BC</sup> | 19,1<sup>AB</sup> | 9,5<sup>A</sup> | 12,1<sup>A</sup> | 9,7<sup>A</sup> | 40,1<sup>AB</sup> | 54,9<sup>BC</sup> | 9,2<sup>A</sup> | 8,3<sup>A</sup> | 7,6<sup>A</sup> | 4,1<sup>A</sup> |
| G1P | 0,4<sup>C</sup> | 0,3<sup>ABC</sup> | 0,1<sup>A</sup> | 0,2<sup>AB</sup> | 0,2<sup>ABC</sup> | 0,2<sup>AB</sup> | 0,3<sup>ABC</sup> | 0,3<sup>BC</sup> | 0,1<sup>A</sup> | 0,2<sup>AB</sup> | 0,2<sup>A</sup> | 0,2<sup>AB</sup> |
| G6P | 3,6<sup>BCDE</sup> | 2,6<sup>ABC</sup> | 2,2<sup>A</sup> | 2,5<sup>AB</sup> | 3,4<sup>BCDE</sup> | 4,5<sup>E</sup> | 4,1<sup>DE</sup> | 3,1<sup>ABCD</sup> | 2,9<sup>ABCD</sup> | 3,3<sup>ABCDE</sup> | 3,8<sup>CDE</sup> | 4,3<sup>DE</sup> |
| Glucose | 39,8<sup>B</sup> | 23,2<sup>AB</sup> | 6,3<sup>A</sup> | 4,2<sup>A</sup> | 4,4<sup>AB</sup> | 4,2<sup>A</sup> | 21,2<sup>AB</sup> | 21,8<sup>AB</sup> | 3,7<sup>A</sup> | 4,2<sup>A</sup> | 4,1<sup>A</sup> | 4,3<sup>A</sup> |
| Starch | 266,2<sup>A</sup> | 313,6<sup>ABC</sup> | 386,6<sup>ABCD</sup> | 410,2<sup>BCD</sup> | 457,4<sup>D</sup> | 442,7<sup>CD</sup> | 268,3<sup>A</sup> | 264,2<sup>A</sup> | 305,8<sup>AB</sup> | 423,3<sup>BCD</sup> | 449,1<sup>D</sup> | 434<sup>BCD</sup> |
| Sucrose | 92,1<sup>ABC</sup> | 67,0<sup>ABC</sup> | 75,5<sup>AB</sup> | 108,3<sup>BCD</sup> | 127,8<sup>CDE</sup> | 160,2<sup>E</sup> | 98,8<sup>ABC</sup> | 68,6<sup>A</sup> | 108,3<sup>BCD</sup> | 141,6<sup>DE</sup> | 156,1<sup>E</sup> | 148,7<sup>DE</sup> |

The accumulation of starch in the pulp cells started very fast and was the highest between 0 and 2 WAE (Figure 4). The balance between the synthesis and the breakdown was clearly in favour of starch synthesis breakdown during the first 8-10 WAE resulting in a net increase in starch content (Figure 4). During the net starch accumulation period, plantain fruit accumulated up to 48% (DW) of starch (Table 2).

**FIGURE 4:**
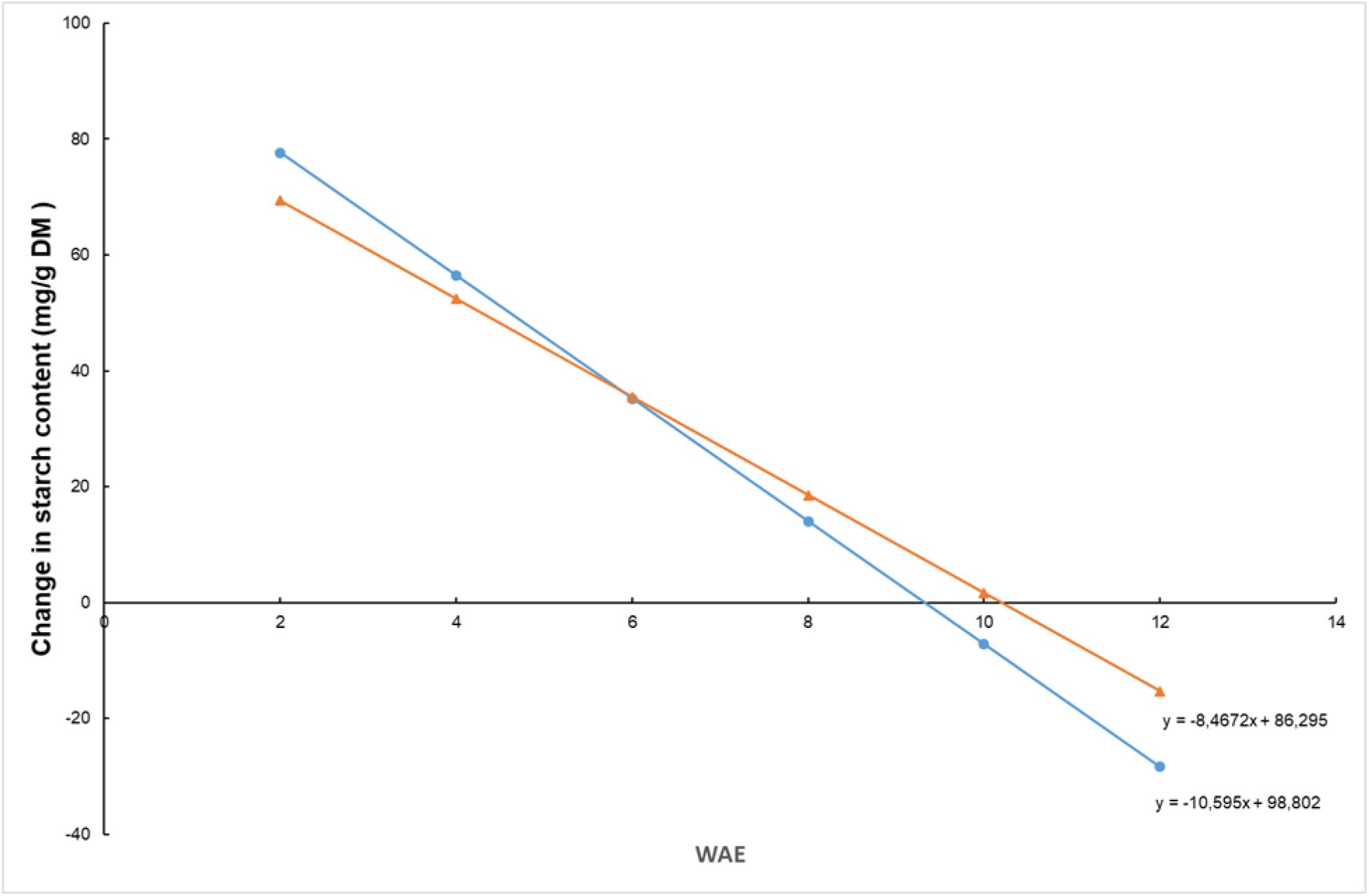
Changes in starch (derivative quadratic regression model). Samples harvested at 2, 4, 6, 8, 10 and 12 WAE. n =4-5 Agbaba (blue); Obino l’ewai (red) Net starch breakdown and so the end of maturation is estimated to take place at 9.3 and 10.2 WAE for Agbaba and Obino l’ewai, respectively.

#### 3.2.1 Starch synthesis

Next to the high abundance of fructose and glucose-6-phosphate (Table 2), a high abundance of a glucose-6-phosphate translocator (6), phosphoglucomutase (5) the glucose-1-phosphate adenylyltransferase (2) and plastidic fructokinase (12) was observed (Table 3, Figure 6A). We did identify a so far uncharacterized sugar translocator (11) (Ma10_p26490) that has an almost perfect correlation (p<0.0001, R=0.99) with SuSy (1) (Table 3, Figure 5).).

**FIGURE 5:**
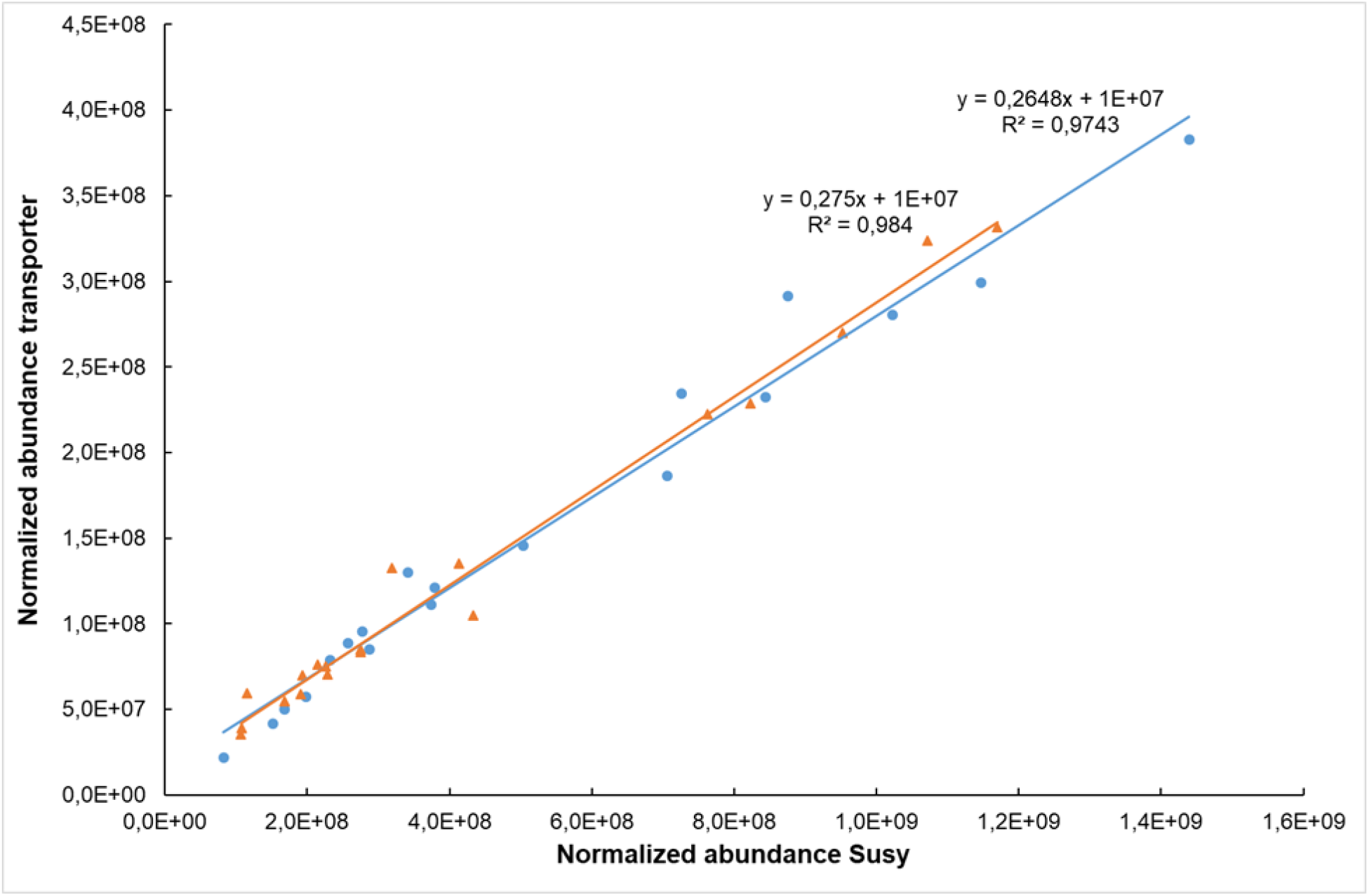
Correlation between plastidic membrane transporter (11, Ma10_p26490) and SuSy (1, Ma08_p23180) abundances. Samples have been harvested at 6, 8, 10 and 12 WAE. Agbaba (blue); Obino l’ewai (red).

**FIGURE 6A:**
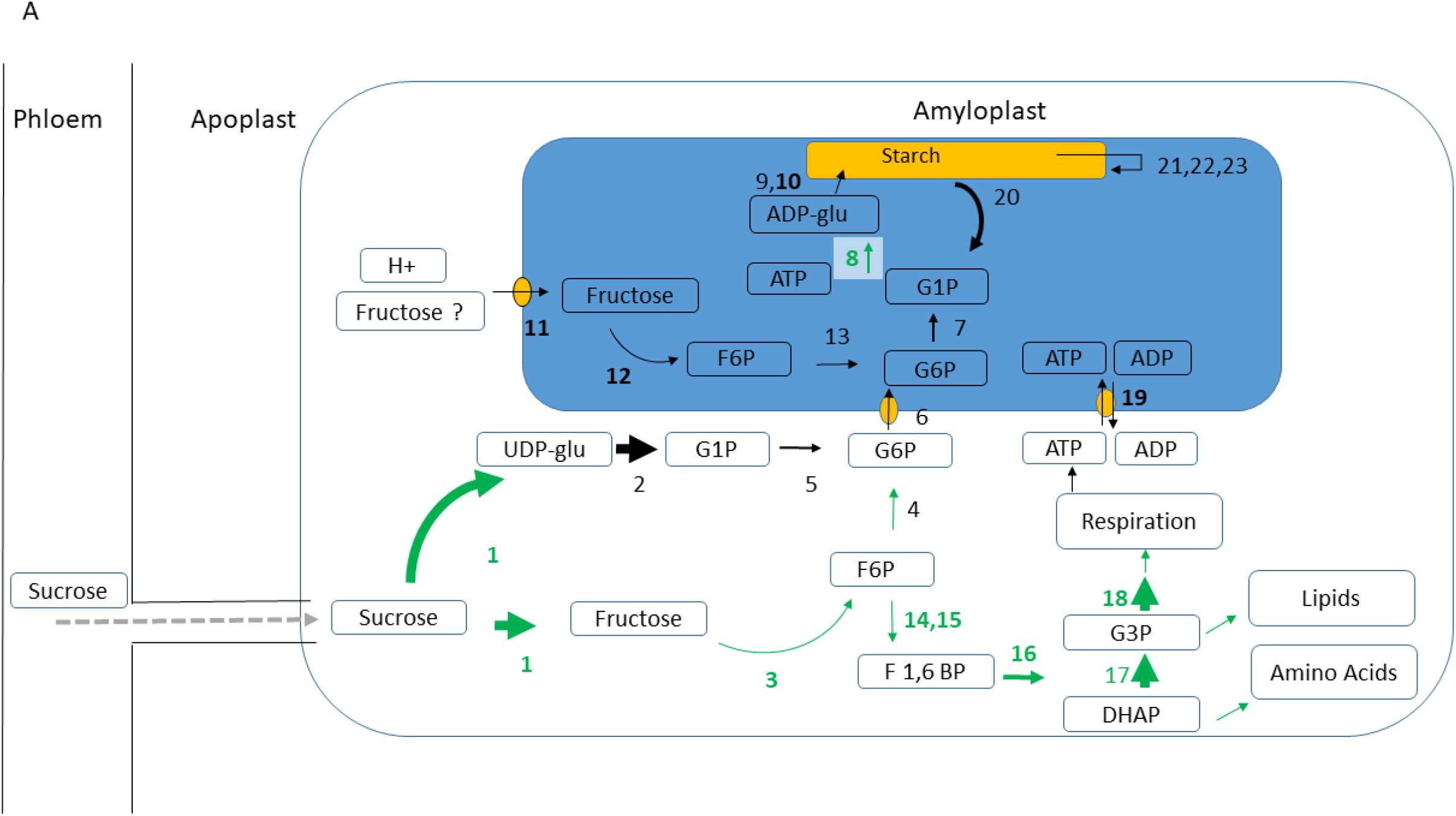
Net starch synthesis at 6 WAE based on proteomic data in two plantain varieties. Enzyme numbers in bold are significantly higher abundant at 6WAE (Table 3). The net direction of the flux is indicated by an arrow. Enzymes and arrows in green have been confirmed by the calculated fluxes (average of two cultivars). The size of the arrow indicates the protein abundance (EMPAI). Grey arrows indicate unidentified or unsure proteins.

**Table 3:**
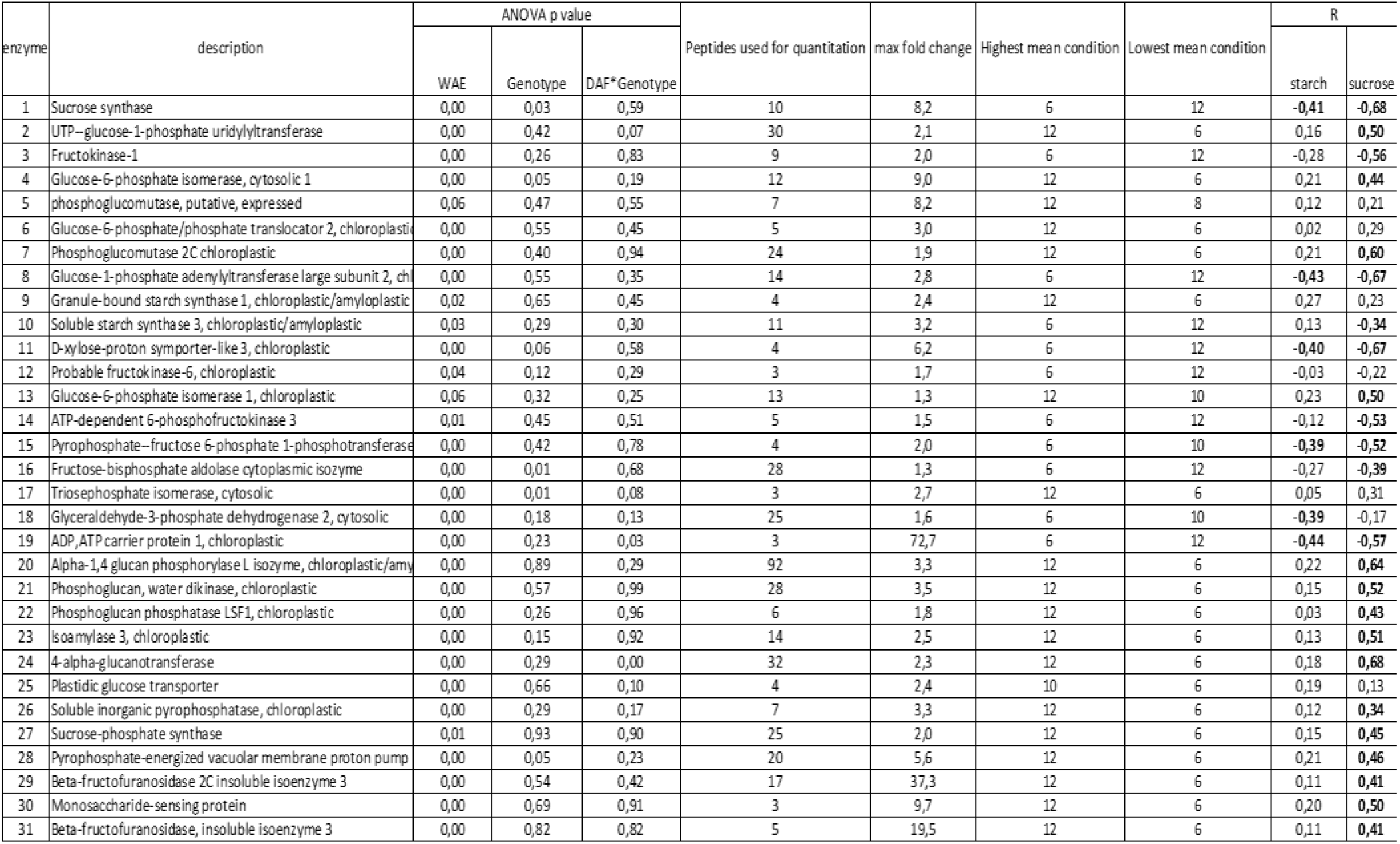
Proteins linked to starch and sugar metabolism and correlations to starch and sucrose in plantain banana pulp. This table only displays the ANOVA p-values of the protein paralog with the lowest value. The full list of significant protein paralogs can be seen in Table S1. WAE: Weeks After Emergence R: Pearson correlation coefficient. Correlations in bold are statistically significant p< 0.05. This table is non redundant and only displays the most significant protein paralogs. The full list can be seen in Table S1

Following the uptake of glucose-6-P into the pulp amyloplast, starch synthesis starts via the concerted action of phosphoglucomutase (7), glucose-1-phosphate adenylyltransferases (AGPase) (8) and the starch polymerizing reactions (9, 10) (Table 3, Figure 6A). In case of fructose, the action of fructokinase (12) and glucose-6-phosphate isomerase (13) are required (Table 3, Figure 6A). The soluble starch synthase (10) decreased in abundance during the further development while granule bound starch synthase raised in abundance (9) (Table 3). Amyloplasts have to import ATP coming from respiration via the cytosol through an ATP/ADP transport protein (19). This enzyme is highly abundant when starch synthesis is high (Table 3, Figure 6A).

Beyond their role as intermediates in the conversion of sucrose to starch, hexose phosphates also serve as substrates for glycolysis and the oxidative pentose phosphate pathway. The significant correlation to starch from pyrophosphate-fructose 6-phosphate 1-phosphotransferase (15) and glyceraldehyde-3-phosphate dehydrogenase (18) (Table 3) is probably due to their function in the glycolysis. Whereas in chloroplasts the ATP necessary for starch synthesis is provided through photosynthesis, in pulp the amyloplasts have to import ATP coming from respiration via the cytosol through an ATP/ADP transport protein (19). This enzyme is highly abundant when starch synthesis is high (Table 3, Figure 6A).

#### 3.2.2 Starch breakdown

The enzymes phosphoglucan water dikinase (PWD) (21), phosphorylase (20), phosphoglucan phosphatases (22), α-1,6-glucosidase starch debranching enzyme (DBE) (23) and 4-α-glucanotransferase disproportionating enzymes (DPE) (24) and the tranporters glucose-6P transporter (6) and the plastidic glucose transporter (25) increased significantly in abundance over time (Table 3, Figure 6B).

**FIGURE 6B:**
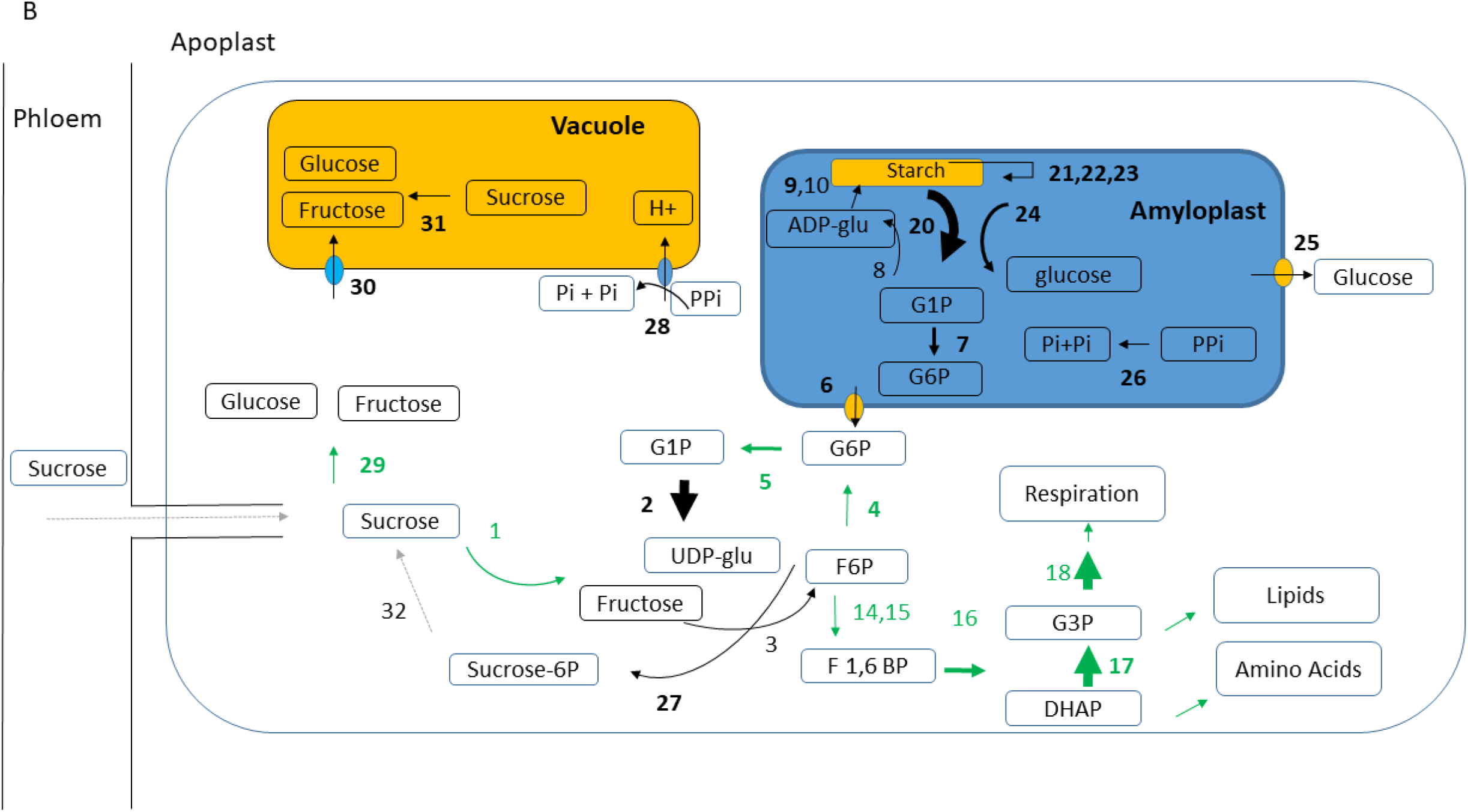
Net starch breakdown at 12WAE. Enzymes in bold are significantly higher abundant at 12WAE (Table 3). The net direction of the flux is indicated by an arrow. Enzymes and arrows in green have been confirmed by the calculated fluxes (average of two cultivars). The size of the arrow indicates the protein abundance (EMPAI). Grey arrows indicate unknown or unsure proteins. 1: Sucrose synthase, 2: UTP-glucose-1-phosphate uridylyltransferase, 3: Fructokinase, 4: Glucose-6-phosphate isomerase, cytosolic, 5: Phosphoglucomutase, 6: Glucose-6-phosphate/phosphate translocator, chloroplastic, 7: Phosphoglucomutase, chloroplastic, 8: Glucose-1-phosphate adenylyltransferase large subunit 2, chloroplastic, 9: Granule-bound starch synthase, chloroplastic/amyloplastic, 10: Soluble starch synthase, chloroplastic/amyloplastic, 11: D-xylose-proton symporter-like 3, chloroplastic, 12: fructokinase, 13: Glucose-6-phosphate isomerase 14: ATP-dependent 6-phosphofructokinase, 15: Pyrophosphate--fructose 6-phosphate 1-phosphotransferase subunit beta, 16: Fructose-bisphosphate aldolase, 17: Triosephosphate isomerase, cytosolic, 18: Glyceraldehyde-3-phosphate dehydrogenase, cytosolic, 19: ADP, ATP carrier protein, chloroplastic, 20: Alpha-1,4 glucan phosphorylase L isozyme, chloroplastic/amyloplastic, 21: Phosphoglucan, water dikinase, chloroplastic, 22: Phosphoglucan phosphatase LSF1, chloroplastic, 23: Isoamylase 3, chloroplastic, 24: 4-alpha-glucanotransferase disproportioning enzyme, 25: Plastidic glucose transporter, 26: Soluble inorganic pyrophosphatase, chloroplastic, 27: Sucrose-phosphate synthase, 28: Pyrophosphate-energized vacuolar membrane proton pump, 29: Invertase, 30: Monosaccharide-sensing protein; 31: Invertase; 32: Sucrose-phosphatase (identification unsure, only 1 peptide)

### 3.3 Sucrose synthesis

The concentration of sucrose significantly increases with time (Table 2). The enzymes with the highest correlation to sucrose were 4-α-glucanotransferase Disproportionating enzymes (DPE) (24), Alpha-1,4 glucan phosphorylase (20) and Phosphoglucomutase, chloroplastic (7) (Table 3). UTP-glucose-1-phosphate uridylyltransferase (2) was one of the most abundant proteins in pulp and its abundance increased with time (Table 3, Figure 6). The production of UGP-glucose can lead to sucrose synthesis either through SuSy (1), which is still abundantly present, or through sucrose-phosphate synthase (27) which had its highest abundance at 12 WAE (Table 3, Figure 6B).

The formed sucrose can then be transported to the vacuole for storage or further processing or can be degraded by invertase (29) and/or SuSy (1) (Table 3, Figure 6B). Invertase (29) Mba10_g13890.1, is only encoded on the B genome and is predicted via the software DeepLoc 1.0 (Almagro Armenteros et al., 2017) to be localized in the cytoplasm. The cytoplasmatic homologue coded on the acuminata genome is most probably not or expressed in a very low level since we did not find a confident specific spectrum. Part of the metabolized sucrose is most likely also transported to the vacuole since we have identified a monosaccharide transporter (30) (Ma04_p22640.1;Mba04_g23280.1) that has the highest abundance at 12 WAE (Table 3, Figure 6B). DeepLoc predicts the membrane protein to the plasma membrane with a likelihood of 0.49 and to the vacuole with a likelihood of 0.35. Since no cell wall invertase has been identified and since we did identify invertase in the cytoplasm, we assume that the monosaccharide transporter (30) is located in the vacuolar membrane (Figure 6B). Also the upregulation of the vacuolar pyrophosphate energized proton pump (28) (Ma07_p22370.1) (Table 3) facilitates the transport of sugars across the vacuolar membrane. We have observed an increased abundance of soluble inorganic pyrophosphatase, in the amyloplast (26) and at the vacuole (28) that coincides with the decrease in starch synthesis and increase in sugars. (Tables 2, 3, Figure 6B).

### 3.4 Cultivar specific ripening

Proteins involved in ascorbate synthesis (GDP-mannose 3,5-epimerase) and anti-oxidant defense (ascorbate peroxidase, Monodehydroascorbate reductase) had their highest abundance at 12 WAE (Table 4). The increase in sucrose production in time is significantly correlated to 1-aminocyclopropane-1-carboxylate oxidase (ACO) (Table 4). We see a cultivar specific interaction between cultivar and WAE meaning that the abundance of ACO changes differently over time in both cultivars (Table 4). A pectinesterase related protein and a lichenase were correlated to both sugar and ACO (Table 4). We did find a GLP (Germin-like protein 12-1) protein that has a significant correlation with ACO (Table 4). A prosite scan shows that the protein has a Fe(^2+^) 2-oxoglutarate dioxygenase domain profile. The protein had moreover also a cultivar specific pattern associated to the earlier ripening Agbagba cultivar (Table 4). We observed an excellent correlation between ACO and a sorbitol dehydrogenase, which catabolize sorbitol into fructose (Fru) and glucose (Glu). Also here the cultivar Agbagba had a significantly earlier response than Obino ‘l Ewai (Table 4).

**Table 4:** Proteins linked to Ethylene response and correlations to ACO in plantain banana pulp WAE: Weeks After Emergence R: Pearson correlation coefficient. Correlations in bold are statistically significant p< 0.05. This table only displays the ANOVA p-values of the protein paralog with the lowest value. The full list of significant protein paralogs can be seen in Table S1.

| Description | ANOVA p value |  |  | Peptides used for quantitation | max fold change | WAE |  | R |  |
| --- | --- | --- | --- | --- | --- | --- | --- | --- | --- |
|  | WAE | Genotype | WAE*Genotype |  |  | Highest mean condition | Lowest mean condition | sucrose | ACO |
| 1-aminocyclopropane-1-carboxylate oxidase | 0,00 | 0,30 | 0,03 | 39 | 19,1 | 12 | 6 | 0,60 | 1,00 |
| Sorbitol dehydrogenase | 0,00 | 0,42 | 0,02 | 13 | 3,4 | 12 | 6 | 0,63 | 0,91 |
| Germin-like protein 12-1 | 0,00 | 0,06 | 0,00 | 5 | 123,8 | 12 | 6 | 0,58 | 0,88 |
| Pectinesterase/pectinesterase inhibitor PPE8B | 0,00 | 0,34 | 0,01 | 18 | 12,8 | 12 | 6 | 0,59 | 0,84 |
| Putative Pectinesterase | 0,00 | 0,86 | 0,39 | 11 | 7,1 | 12 | 6 | 0,57 | 0,82 |
| L-ascorbate peroxidase, cytosolic | 0,00 | 0,07 | 0,00 | 14 | 2,5 | 12 | 6 | 0,57 | 0,82 |
| GDP-mannose 3,5-epimerase 1 | 0,00 | 0,21 | 0,56 | 13 | 1,9 | 12 | 6 | 0,50 | 0,82 |
| Probable L-ascorbate peroxidase 7, chloroplastic | 0,00 | 0,12 | 0,87 | 7 | 3,4 | 12 | 6 | 0,50 | 0,75 |
| Lichenase | 0,00 | 0,23 | 0,00 | 20 | 46,1 | 12 | 6 | 0,56 | 0,73 |

### 3.5 The global flux decreased throughout fruit development

The measured concentrations of the biomass and the accumulated metabolites in the pulp, determined at 2, 4, 6, 8, 10 and 12 WAE (Table 2) were fitted to calculate the corresponding fluxes used as constraints in the metabolic model. The estimated fluxes showed the highest activity for fluxes involved in respiration, glycolysis and TCA cycle (Figure S3). At the early stage of development (Figure 7, 2 WAE) those fluxes had their heighest activity and a global decrease throughout fruit development was noticed, in agreement with metabolic fluxes described on tomato fruit (Colombié et al., 2015). This decrease in flux activity throughout fruit development was similar in both cultivars. We assessed that a high respiration is associated with the cell division associated with the first growth phase followed by a global decrease in flux activity during the second and third-growth phase, where only elongation takes place. No increase in respiration was detected at the end of the maturation, probably because the burst did not take place yet in the investigated fruits.

**Figure 7:**
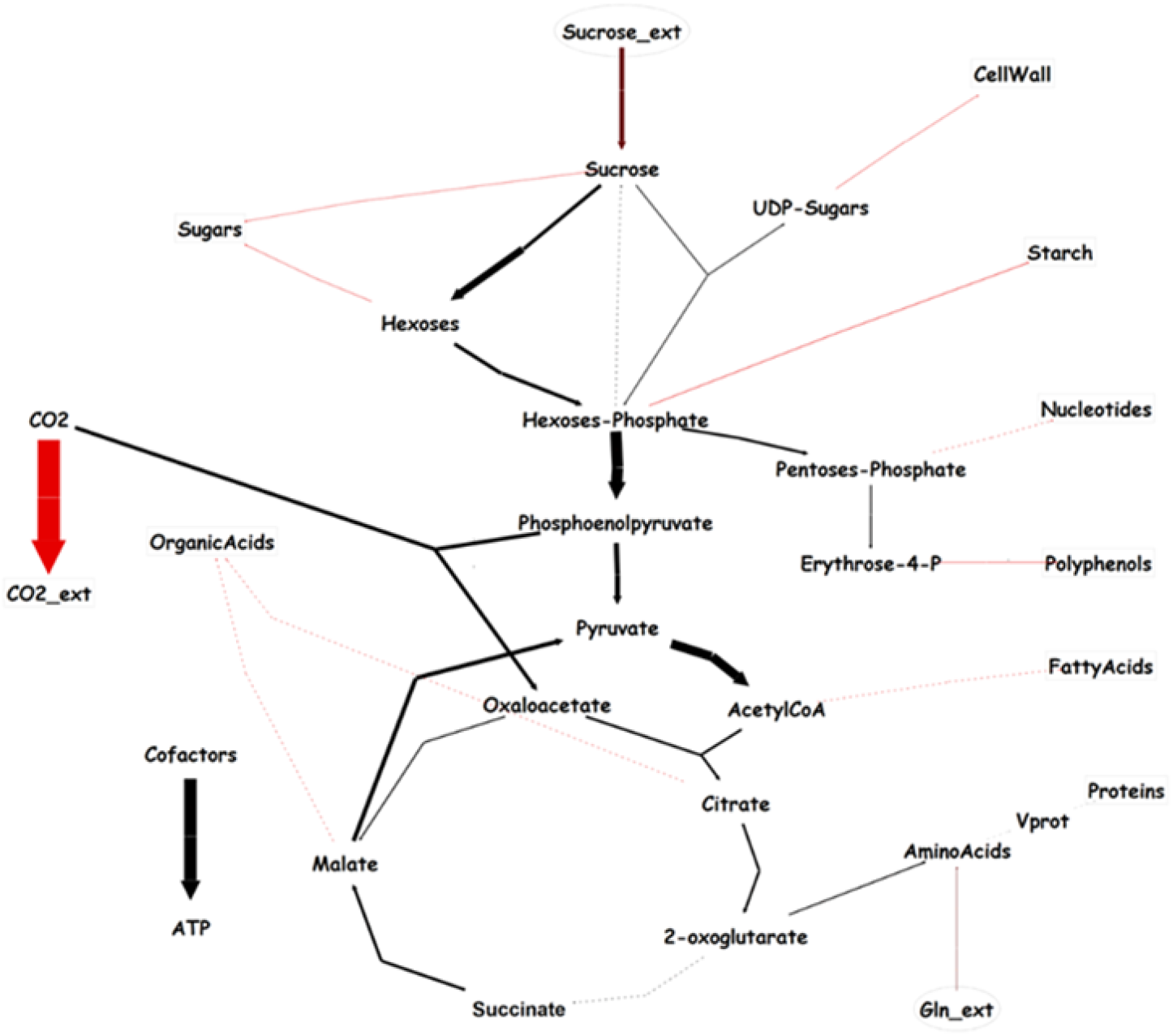
Simplified flux map, based on constrain-based modelling for Agbaba plantain cultivar at 2 WAE, showing a high activity for fluxes in glycolysis, TCA cycle and mostly respiration (in red). The same trend was obtained for both cultivars (see Figure S3). The arrow width is proportional to flux intensity.

The flux analysis complemented the proteome data (Figures 6 A and B). Nice concordance was observed for major reactions in starch biosynthesis: sucrose synthase (1), fructokinase (3), glucose-6-phosphate isomerase (4), and glucose-1-phosphate adenylyltransferase (8) (Figure 6A).

For the flux at 12 WAE in the starch degradation pathway (Figure 6B) next to invertase (29) also fluxes through sucrose synthase (1), and glucose-6-phosphate isomerase (4) pointed towards a net cleavage of sucrose. Some uncertainties in flux calculations might be attributed to the assumptions required to solve the model (flux minimization).

## 5 Discussion

### 5.1 Three different fruit growth phases with their particular proteome and metabolic profile

In banana, the growth pattern is cultivar dependent and fertilization influences the growth and the shape of the fruit (Simmonds, 1953). A sigmoid type of growth has been described before in a triploid banana with a B (balbisiana) genome (Awak legor) (Simmonds, 1953). The first period of fast growth is characterized by cell division and cell elongation, while the second one is due to cell elongation only (Ram et al., 1962). The increase in cell number in the initiating region of the pulp has been reported to continue up to about 4 WAE in the cultivar Pisang lilin (a partenocarp AA) (Ram et al., 1962). We did find evidence to confirm the involvement of cell division in growth in our proteomics data. Histones are one of the primary components of chromatin and are synthesized during the S-phase. The speed of DNA replication is depending on the rate of histone biosynthesis (Ma et al., 2015).

Banana pulp tissue is a starch synthesizing sink tissue that needs to get all its energy from the sucrose unloaded from the phloem and from starch degradation. From tomato, it is known that the fruit growth consists of two phases: (i) a period of rapid fruit growth where sucrose synthase is determining the sink strength, and (ii) a phase after rapid growth has ceased, where invertase takes over (Nguyen-Quoc and Foyer, 2001). Based on the observed abundance pattern of Sucrose Synthase (SuSy) and invertase, we hypothesize that the first fast growth phase is completely dominated by Sucrose Synthase (SuSy), while the second fast growth phase is dominated by invertase. The abundance of invertase showed an excellent correlation with the growth rate (Figure 3).

### 5.2 Starch synthesis: cytosolic glucose -6 phosphate and fructose are important sources for starch synthesis

Because starch synthesis in pulp is confined to amyloplasts, it relies entirely on translocation of metabolites from the cytosol through the amyloplast envelope. The form in which carbon enters the amyloplast has long been a matter of debate (Hofius and Börnke, 2007). The triose phosphate transporter from chloroplasts is a perfectly annotated and studied transporter in the plastid envelope of many plants. However, there is discussion as to whether the genes are expressed in non-green tissue (Tobias et al., 1992; Neuhaus and Emes, 2000). In potato it is clear that triose phosphate is not the substrate taken up to support starch synthesis (Hofius and Börnke, 2007). Our data also point into the same direction since we were not able to identify a triose phosphate transporter protein in plantain pulp. Amyloplasts of tubers or fruits are also normally not able to generate hexose phosphates from C3 compounds due to the absence of fructose 1,6-bisphosphatase activity (Nguyen-Quoc and Foyer, 2001; Hofius and Börnke, 2007). They rely on the import of cytosolically generated hexose phosphates as the source of carbon for starch biosynthesis (Entwistle and Rees, 1988; Hofius and Börnke, 2007). This seems also to be the case here in plantain since we were not able to identify a fructose 1,6-bisphosphatase protein during the period of investigation. The enzyme does seem active though in non-photosynthetic tissues where it controls the rate of F6P production in the gluconeogenetic pathway (Hofius and Börnke, 2007). We did identify the enzyme though in low quantities in our previous analysis were we analyzed ripening detached fruits (Bhuiyan et al., 2020). So also in our case it might play a role in the starch breakdown much later when the ripening and sugar synthesis is more advanced. None of the three predicted adenine nucleotide BT1 transporters (Ma10_p26970, Ma07_p09880, Ma06_p06780) that transport ADP-glucose across the plastid membrane was identified in the present study. Therefore, it is also unlikely that ADP-glucose is moving across the amyloplast envelope to provide substrates for starch synthesis. We suggest that in plantain banana the cytosolic glucose -6 phosphate is an important direct source of sugar for starch synthesis as it is the case in maize (Tobias et al., 1992). This was confirmed by the high abundance of the glucose-6-phosphate/phosphate translocator (6), phosphoglucomutase (5) and the glucose-1-phosphate adenylyltransferase (2) (Table 3). We did identify a so far uncharacterized sugar translocator (11) (Ma10_p26490) that has an almost perfect correlation (p<0.0001, R=0.99) with SuSy (1) (Table 3, Figure 5). Plastids are able to transport sugars across their membranes (Patzke et al., 2019). However, only two plastidic sugar transporters are well known and described (Weber et al., 2000; Niittylä et al., 2004). These transporters reside in the inner envelope membrane and respectively mediate the export of maltose and glucose (Cordenunsi-Lysenko et al., 2019). Considering our observed tight correlation with SuSy, we hypothesize that the Ma10_p26490.1 transporter transports fructose across the amyloplast membrane. The abundance pattern of the plastidic fructokinase (12) corroborates this hypothesis (Table 3, Figure 6A). Since only very few reports are available on plastid fructose/glucose/sucrose H transporters (Patzke et al., 2019), more studies are needed to confirm our hypothesis and confirm its physiological role in starch synthesis.

The soluble starch synthase (10) decreased in abundance during the further development while granule bound starch synthase raised in abundance (9) (Table 3). This abundance pattern suggests that during the early starch synthesis, soluble starch synthase is more important. The fact that the polymerizing reactions of starch synthesis are not dominant in the control of starch accumulation has to do with the balance between sink strength, starch synthesis and starch breakdown and has been observed before (Tetlow et al., 2004). So based on the ANOVA analysis and the correlations, the main drivers of starch synthesis in plantain pulp seem to be Sucrose Synthase (1), Glucose-1-phosphate adenylyltransferase (ADP-glucose pyrophosphorylase (AGPase)) (8), ADP, ATP carrier protein (19) and the so far uncharacterized membrane sugar transporter (11) (Figure 6A, Table 3).

### 5.3 Starch breakdown: phosphoglucan water dikinase, alpha-1,4 glucan phosphorylase, phosphoglucan phosphatase, isoamylase and 4-alpha-glucanotransferase initiate breakdown

The starch-to-sucrose metabolism has been extensively studied in model systems in the context of energy sources for plant growth and development (Streb and Zeeman, 2012). However, the starch breakdown in fleshy fruits such as bananas is less understood (Cordenunsi-Lysenko et al., 2019). All the genes involved in starch breakdown have been mapped on the banana genome (Xiao et al., 2018). Based on what is known from Arabidopsis, it was hypothesized that in banana starch-phosphorylating enzymes, termed glucan water dikinase (GWD), phosphorylate the C6 position and the phosphoglucan water dikinase (PWD) phosphorylate the C3 position of the glycosyl residues in starch (Cordenunsi-Lysenko et al., 2019). The role of phosphorylases including GWD and PWD in starch breakdown during banana ripening is less understood, but phosphorylation at the C3 and C6 position of the glucosyl residues in the starch of freshly harvested unripe bananas has already been found, as well as the presence of PWD and GWD (Cordenunsi-Lysenko et al., 2019). The steric hindrance of these phosphorylated groups alters the organization of the granule and it has been hypothesized that PWD acts downstream of GWD and that the induced phosphorylation of banana starch favors granule hydration and phase transition from the crystalline state to the soluble state (Cordenunsi-Lysenko et al., 2019). Our data confirm that dikinases play a role in early starch breakdown but not that PWD would act downstream of GWD. We have identified the sole PWD protein present in the banana genome (21) (Ma09_p07100.1;Mba09_g06570.1) as being present at the early stage of starch breakdown process and being significantly upregulated (Table 3) while none of the two GWD proteins could be detected. We did identify GWD1 in our previous study during the ripening of detached plantain fruits (Bhuiyan et al., 2020) and also Xiao and coworkers identified GWD1 in ripening detached fruits as being expressed at the late ripening stages (Xiao et al., 2018).

Phosphorolytic cleavage seems to be one of the first starch breakdown reactions. This hypothesis is corroborated by the abundance profiles of phosphorylase (20) and from the glucose-6P transporter (6) (Table 3, Figure 6B). The increase in abundance and activity of phosphorylase was also observed when investigating phosphorylase during maturation and ripening (Da Mota et al., 2002). Also other enzymes appear to contribute to the early degradation of starch. Phosphoglucan phosphatases (22), α-1,6-glucosidase starch debranching enzyme (DBE) (23) and 4-α-glucanotransferase Disproportionating enzymes (DPE) (24) increase significantly in abundance (Table 3, Figure 6B). We also observed the increased abundance of the plastidic glucose transporter (25) (Table 3, Figure 6B), while the Maltose transporter, Maltose Excess Protein transporter was not detected. Since also neither alpha nor beta-amylases were detected at this early stage of ripening, we hypothesize that they act later in the ripening process. While investigating detached ripening fruits, we found that plastidic alpha amylase acts before beta amylase (Bhuiyan et al., 2020). This was also found by (Purgatto et al., 2001). Beta amylase is essential to complete the breakdown and its upregulation was reported to be correlated to a decrease in starch during fruit ripening (Purgatto et al., 2001; Bhuiyan et al., 2020)

### 5.4 Sucrose synthesis: competition between vacuolar storage and recycling sucrose for growth and starch resynthesis

Starch breakdown products G1P and glucose are produced which can be metabolized further. The cytoplasmic G1P has been proven to flow to the production of UGP-glucose (Figure 6B). UGP-glucose can lead to sucrose synthesis either through SuSy (1), which is still abundantly present, or through sucrose-phosphate synthase (27) which has its highest abundance at 12 WAE (Table 3, Figure 6B). We did not confidently identify sucrose phosphatase at this early stage of ripening. Only one peptide was found with low confidence. The reason for the low confidence is probably the low abundance of the enzyme. We did confidently identify sucrose phosphatase in our study of detached ripening fruits; it proved to be low abundant and was significantly upregulated in the very late ripening stages (Bhuiyan et al., 2020). The formed sucrose can then be transported to the vacuole for storage or further processing or can be degraded by invertase (29) and/or SuSy (1) (Table 3, Figure 6B). Most banana production, both of dessert and cooking types, is based on triploid cultivars. Banana cultivars are natural combinations of different A (acuminata) and B (balbisiana) genomes and have been fixed over hundreds of years of human selection. Plantain is an allopolyploid crop with an AAB genome (Carreel et al., 2002). Invertase (29) Mba10_g13890.1, is only encoded on the B genome. The cytoplasmatic homologue coded on the acuminata genome is most probably not or expressed in a very low level since we did not find a confident specific spectrum. We have shown before that invertase is more abundant in plantain compared with a Cavendish type (Bhuiyan et al., 2020). A higher invertase activity in cooking bananas has already been associated with a changed sucrose/(glucose + fructose) ratio (Fils-Lycaon et al., 2011). The breakdown of sucrose in the cytoplasm by invertase would enable to flow back to starch synthesis and glycolysis to support further growth as discussed above (Figure 3). Plantains are indeed a lot bigger than dessert bananas and contain much more starch. Part of the metabolized sucrose is most likely also transported to the vacuole since we have identified a monosaccharide transporter (30) (Ma04_p22640.1;Mba04_g23280.1) that has the highest abundance at 12 WAE (Table 3, Figure 6B). Also the upregulation of the vacuolar pyrophosphate energized proton pump (28) (Ma07_p22370.1) (Table 3) facilitates the transport of sugars across the vacuolar membrane (Maeshima, 2000). Alterations in PPi metabolism have a strong effect on sugar metabolism in which higher PPi levels increase starch accumulation and decrease the level of sucrose. Decreased PPi levels have been associated with lower starch biosynthetic rates (Osorio et al., 2013). The overexpression of a pyrophosphatase in tomato resulted in an increase in the major sugars, a decrease in starch and an increase in vitamin C (ascorbic acid) (Osorio et al., 2013). We have observed an increased abundance of soluble inorganic pyrophosphatase, in the amyloplast (26) and at the vacuole (28) that coincides with the decrease in starch synthesis and increase in sugars. (Tables 2, 3, Figure 6B). Indeed also proteins involved in ascorbate synthesis (GDP-mannose 3,5-epimerase) and anti-oxidant defense (ascorbate peroxidase, Monodehydroascorbate reductase) were higher abundant at 12 WAE (Table 4). Ascorbic acid is also a cofactor of 1-aminocyclopropane-1-carboxylic acid oxidase (ACO) that catalyzes the final step in the biosynthesis of the plant hormone ethylene (Smith et al., 1992).

### 5.5 Cultivar specific ethylene biosynthesis and auxin scavenging

Climacteric fruits show a dramatic increase in the rate of respiration during ripening and this is referred to as the climacteric rise (Paul et al., 2012). The rise in respiration is logarithmic and occurs either simultaneously with the rise in ethylene production or it follows soon afterwards (Burg, 1962). However, this large change in the magnitude of ethylene production can be misleading. The important point is when the tissue becomes more sensitive to ethylene and internal concentration reaches a threshold concentration required to induce biological responses (Paul et al., 2012). Thus, ethylene plays a major role in the ripening process of climacteric fruits. Climacteric fruits can ripen fully if they are harvested at completion of their growth period. We finish our analysis at this point since this is the point that plantains are harvested and consumed. The increase in sucrose production in time is significantly correlated to 1-aminocyclopropane-1-carboxylate oxidase (ACO) (Table 4). ACO is the enzyme that produces ethylene. It is well-known that banana is a climacteric fruit and so that ripening and net sugar synthesis starts upon ethylene production (Cordenunsi and Lajolo, 1995; do Nascimento et al., 2000; Cordenunsi-Lysenko et al., 2019). Banana has two interconnected feedback loops (Lü et al., 2018). The first one is a positive feedback loop dependent on NAC transcription factors, while the second one is controlled by MADS transcription factors and is able to maintain the ethylene synthesis even when the first loop is blocked. It has been shown that banana ACO has a NAC motif in the promoter sequence (Lü et al., 2018). It has been illustrated that ripening is a highly coordinated process regulated at the transcript level (Kuang et al., 2021). We see a cultivar specific interaction between cultivar and WAE meaning that the abundance of ACO changes differently over time in both cultivars (Table 4). The disappearance of the large stock of starch in favour of the accumulation of soluble sugars has also already been proven to contribute to pulp softening (Shiga et al., 2011). A pectinesterase related protein and a lichenase are associated to pulp softening (Li et al., 2019; Bhuiyan et al., 2020) and were correlated to both sugar and ACO (Table 4). Proteins with sequence similarity to germins have been identified in various plant species. Those ‘germin-like proteins’ (GLPs) have a global low sequence identity with germins and constitute a large and highly diverse family with diverse functions among them auxin binding (Bernier and Berna, 2001). Two auxin correlated GLPs were isolated in plum that were correlated to the change of levels of autocatalytic ethylene levels and associated ripening (El-Sharkawy et al., 2010). The authors found differential expression in two contrasting cultivars and hypothesized that the differential endogenous auxin levels in the two cultivars change the levels of available ethylene and so the ripening phenotype. We did find a GLP (Germin-like protein 12-1) protein that has a significant correlation with ACO (Table 4). A prosite scan shows that the protein has a Fe(^2+^) 2-oxoglutarate dioxygenase domain profile. A 2-oxoglutarate-dependent-Fe (^2+^) dioxygenase in rice has been shown to convert active auxin (indole acetic acid) into biologically inactive 2-oxoindole-3-acetic acid, supporting a key role in auxin catabolism (Zhao et al., 2013). The protein has moreover also a cultivar specific pattern associated to the earlier ripening Agbagba cultivar (Table 4). We hypothesize that this GLP/2-oxoglutarate dioxygenase would catabolize auxin and hence stimulate ripening. In banana it has been proven that ethylene promotes ripening and auxins delay it (Purgatto et al., 2001; Mainardi et al., 2006; Kuang et al., 2021). Also in papaya the same has been proven (Zhang et al., 2020).

In plum, it has been shown that ethylene was a crucial factor affecting overall sugar metabolism (Farcuh et al., 2018). More specifically, ethylene reduced sucrose catabolism and induced sucrose biosynthesis but inversely, stimulated sorbitol breakdown via increased sorbitol and dehydrogenase decrease sorbitol biosynthesis via decreased sorbitol-6-phosphate-dehydrogenase. Also here, we observed an excellent correlation between ACO and a sorbitol dehydrogenase, which catabolize sorbitol into fructose (Fru) and glucose (Glu). Also here the cultivar Agbagba has a significantly earlier response than Obino ‘l Ewai (Table 4).

## 6 Conclusions

By combining proteomics and flux studies, we gain here unique insights into the order of appearance and dominance of specific enzymes/fluxes involved in starch and sugar synthesis and breakdown. Fluxes give a broader analysis of the metabolism. Despite fluxes are calculated in a non-compartmented network, we showed that proteome data complemented by fluxes can give a satisfactory picture of the dynamics of metabolism during fruit development. The maturation in plantain is completed around 10 WAE, indicated by a net breakdown in starch. The import of G6P into the amyloplast and possibly fructose are the main drivers of starch synthesis. The soluble starch synthase likely plays a more important role in the starch synthesis during the early fruit development while granule bound starch synthase most likely influences the starch at the mature stage. For starch breakdown, mainly DPE and phosphorylase produce the first hexoses for sugar synthesis and amylases come into play at a later stage in ripening. In plantain cytoplasmic invertase seems to play an important role in the breakdown of sucrose to support further growth. The data pointed towards an interplay between auxins and ethylene, controlling the ripening process. Despite the fact that both plantain cultivars are extremely close genetically, we did find significant differences in ripening. The earlier ripening in Agbagba might be related to an earlier induction of the second ethylene system and a bigger scavenging of auxins. This information contributes to a better understanding of fruit development and maturation in banana and more specifically plantains.

## Acknowledgements

The authors would like to thank Kusay Arat for the technical support at SYBIOMA, KU Leuven, Belgium. We acknowledge USAID for the project AID-BFS-G-II-00002-11 Reviving the plantain breeding program at IITA – International Institute for Tropical Agriculture. The authors would furthermore like to thank all donors who supported this work through their contributions to the CGIAR Fund (http://www.cgiar.org/who-we-are/cgiar-fund/fund-donors-2/), and in particular to the CGIAR Research Program on Roots, Tubers and Bananas, and the PHENOME (French ANR-11-INBS-0012) project for funding. The metabolite analyses were performed on Bordeaux Metabolome facility.

The authors have declared no conflict of interest.

